# RNA-catalyzed RNA Ligation within Prebiotically Plausible Model Protocells

**DOI:** 10.1101/2023.02.09.527907

**Authors:** Saurja DasGupta, Stephanie J. Zhang, Merrick P. Smela, Jack W. Szostak

**Author notes:** **Corresponding Author Jack W. Szostak** − Massachusetts General Hospital, Boston, Massachusetts, and Harvard Medical School, Boston, Massachusetts. These authors contributed equally to this work (Saurja DasGupta, Stephanie J. Zhang). Dept. of Chemistry, University of Chicago, Chicago IL 60635.

## Abstract

Demonstrating RNA catalysis within prebiotically relevant models of primordial cells (protocells) remains a challenge in Origins of life research. Fatty acid vesicles encapsulating genomic and catalytic RNAs (ribozymes) are attractive models for protocells; however, RNA catalysis has largely been incompatible with fatty acid vesicles due to their instability in the presence of Mg^2+^ at concentrations required for ribozyme function. Here, we report a ribozyme that catalyzes template-directed RNA ligation at low Mg^2+^ concentrations and thus remains active within stable vesicles. Ribose and adenine, both prebiotically relevant molecules, were found to greatly reduce Mg^2+^-induced RNA leakage from vesicles. When we co-encapsulated the ribozyme, substrate, and template within fatty acid vesicles, we observed efficient RNA-catalyzed RNA ligation upon subsequent addition of Mg^2+^. Our work shows that RNA-catalyzed RNA assembly can occur efficiently within prebiotically plausible fatty acid vesicles and represents a step toward the replication of primordial genomes within self-replicating protocells.

## Introduction

Life is thought to have originated via a series of chemical steps involving a gradual increase in molecular and organizational complexity. In an RNA-centric view of the origin of life, catalytic RNAs or ribozymes emerged from the enzyme-free assembly of oligonucleotides which were themselves replicated via the nonenzymatic polymerization of activated monomers or ligation of activated oligomers. (1-3) Ribozymes capable of accelerating RNA assembly would drive genome replication and generate a diverse collection of functional RNAs, and thus shape the early evolution of life. However, biological evolution would not have been possible in the absence of compartmentalized catalysis. Cellular compartments provide a physical separation between the individual and its surroundings and therefore allow beneficial phenotypes to confer selective advantage to the individual. (1, 4, 5) In the primordial biology of the RNA World, compartments would have been necessary to keep RNA genomes together with their encoded ribozymes, (6) and to limit takeover by parasitic molecules. (7) Once catalytic phenotypes began to provide beneficial effects on cellular stability, growth, and division, Darwinian evolution would have rapidly accelerated the development of increasingly complex living systems. (1)

We are developing experimental models for the major chemical transitions that led to the emergence of the earliest cells (protocells). For example, we have demonstrated nonenzymatic copying of short RNA sequences via the polymerization of monomers activated with the prebiotically relevant 2-aminoimidazole (2AI) moiety. (8) Nonenzymatic ligation of 2AI-activated RNA oligonucleotides is considerably slower than primer extension, (9) which prompted us to identify ribozymes that use 2AI-activated RNA oligonucleotides as substrates for ligation. (10) Since the first ribozymes are likely to have been generated through the nonenzymatic assembly of intrinsically reactive oligonucleotide building blocks, (11, 12) these ‘2AI-ligase’ ribozymes can be considered to be intermediates in the transition between nonenzymatic and RNA-catalyzed RNA assembly.

Given the importance of compartmentalization to biological evolution, life most likely emerged as protocellular entities consisting of simple genomes and enzymes made of RNA, encapsulated within primitive compartments. Membrane-less compartments such as coacervates have been shown to support nonenzymatic RNA assembly and ribozyme catalysis and have also been proposed as models of primordial cells. (13-17) However the requisite components of coacervates that have been reported in the literature, including polydiallyldimethylammonium chloride, polyallylamine, polyaspartic acid, polyarginine, and polylysine, are not prebiotically relevant. In addition, the rapid exchange of RNA molecules between coacervate droplets (18) would make coacervates ill-suited for evolutionary adaptation.

All known life consists of cells bounded by bilayer membranes composed of amphiphilic molecules. Fatty acids, the primary amphiphilic component of diacyl phospholipids, are excellent candidate constituents of primitive cell membranes as they readily self-assemble into vesicles, (19-21) that can encapsulate and retain genetic material. (22) Fatty acids can be synthesized under simulated prebiotic conditions, (23-25) and have been detected in abiotic environments such as carbonaceous meteorites. (26) Fatty acid vesicles can grow in response to increased internal osmotic pressure due to the accumulation of encapsulated RNA (27-29) and multilamellar fatty acid vesicles have been observed to grow into filamentous vesicles that divide under mild agitation. (30, 31) Giant unilamellar vesicles (GUVs) can grow and divide following the addition of fatty acid micelles. (32) Membranes composed of a combination of short chain (<C12) fatty acids, alcohols, and glycerol esters are permeable to RNA building blocks such as nucleoside phosphorimidazolides and to cations such as Mg^2+^ that are essential for both enzymatic and enzyme-free RNA assembly as well as other RNA-catalyzed processes. These properties make fatty acid vesicles attractive for modeling compartmentalized RNA assembly. (33)

To model the initial stages of RNA replication within protocells, our lab previously demonstrated nonenzymatic template-directed primer extension using nucleoside phosphorimidazoles within fatty acid vesicles. (33, 34) As vesicles made of prebiotically relevant fatty acids such as decanoic acid tend to be destabilized at Mg^2+^ concentrations of ∼3 mM and higher, a high concentration of citrate-chelated Mg^2+^ was used in these experiments to supply sufficient Mg^2+^ for RNA assembly, while preventing vesicle destabilization. (33) However, such a high local concentration of citrate is prebiotically unlikely. (1) In principle, polymerase ribozymes could significantly accelerate RNA assembly within protocells; however, these ribozymes, in their current form, work exclusively with nucleoside triphosphates (NTPs) and have high Mg^2+^ concentration requirements (>100 mM) making them incompatible with fatty acid compartments. (35, 36) Furthermore, the high negative charge on NTPs prevents rapid permeation across fatty acid membranes at room temperature, precluding RNA copying within prebiotic compartments using NTPs as external ‘nutrients’. (37) RNA-catalyzed RNA assembly using 2AI-activated monomers supplied from the extracellular environment could provide a means for protocellular genome replication and RNA synthesis-driven protocellular growth and division. (27, 30) As polymerase ribozymes that use NTP substrates have been iteratively evolved from ribozymes that catalyze the ligation of 5′ triphosphorylated oligoribonucleotides, (1) we reasoned that the first step toward constructing a protocell that supports ribozyme-polymerase catalyzed RNA replication would be to demonstrate RNA-catalyzed RNA ligation within such compartments. Here, we show that RNA-catalyzed RNA assembly within prebiotically plausible fatty acid protocells is possible at low concentrations of Mg^2+^ using a ligase ribozyme that uses 2-aminoimidazole (2AI)-activated RNA oligonucleotide substrates.

## Results and Discussion

### A ligase ribozyme that functions at low Mg^2+^ concentrations

We previously isolated several classes of ligase ribozymes that catalyze the formation of a 3′-5′ phosphodiester bond between an RNA primer (covalently attached to the catalytic domain of the ribozyme via a polyuridine linker) and a 2-aminoimidazole (2AI)-activated RNA oligonucleotide, aligned on an external template. (10) In that work, we characterized the most abundant sequences (RS1-RS10) from each of the ten most abundant ribozyme classes isolated from *in vitro* selection, and found that truncated versions of four (RS1, RS5, RS7, RS8) of the ten ligases retained function. The catalytic domains of these truncated ribozymes are 56 nt-long, and therefore more accessible to nonenzymatic RNA assembly than the longer parent sequences. (10) Because the full-length RS7 was found to exhibit lower ligation yields compared to the other three ribozymes (∼20% vs >50% in 2 h at 10 mM Mg^2+^), we removed it from further consideration. As the 2AI-ligase ribozymes were selected under modest (5-20 mM) concentrations of Mg^2+^, we wondered if any of these full-length ribozymes would retain activity in an even lower Mg^2+^ concentration regime. We tested ligation of a 16 nt 2AI-activated substrate by full-length ribozymes, RS1, RS5, and RS8 in the presence of 1-5 mM Mg^2+^ (Supplementary Figure S1). At Mg^2+^ concentrations below 3 mM, RS1 and RS5 exhibited poor ligation (<10% ligation after 4 h), while RS8 exhibited >50% ligation (Supplementary Figure S1A-C and Supplementary Figure S2). RS1 exhibited ligation yields of ∼14% and 28% at 4 mM and 5 mM Mg^2+^, respectively, while RS5 ligated to 18% and 25% at 4 mM and 5 mM Mg^2+^, respectively. RS8 was significantly more efficient than both RS1 and RS5 under these conditions with ∼60% ligation after 3 h (Supplementary Figure S1D, E and Supplementary Figure S2).

Since the full length RS8 sequence was the most efficient ribozyme at low [Mg^2+^], we tested its truncated version, RS8_trunc, for ligation activity at low [Mg^2+^] (Figure 1A). Ligation was detectable at concentrations as low as 0.6 mM and ligation rate increased with an increase in [Mg^2+^], saturating at 2 mM (Figure 1B and Supplementary Figure S3). Ligation yields increased from ∼15% in 0.6 mM to ∼ 40% in 2-4 mM Mg^2+^ after 3 h (Figure 1C). These assays were performed in the presence of 250 mM Na^+^, the same conditions used in the *in vitro* selection experiment from which these 2AI-ligase ribozymes were isolated. However, high concentrations of monovalent cations can be detrimental to the stability of vesicles composed of prebiotically relevant fatty acids. (38) Therefore, we tested ligation at various concentrations of Na^+^ in the background of 3 mM Mg^2+^, which revealed a modest dependence of ribozyme activity on Na^+^ and, more importantly, significant catalytic ligation in the absence of Na^+^ (Supplementary Figure S4). Therefore, all subsequent ligation reactions were performed without Na^+^. The ability of RS8_trunc to catalyze ligation at low [Mg^2+^] and in the absence of Na^+^, conditions that are compatible with the stability of fatty acid compartments, suggested that it might be possible to constitute RNA-catalyzed RNA ligation within prebiotically relevant model protocells.

**Figure 1.**
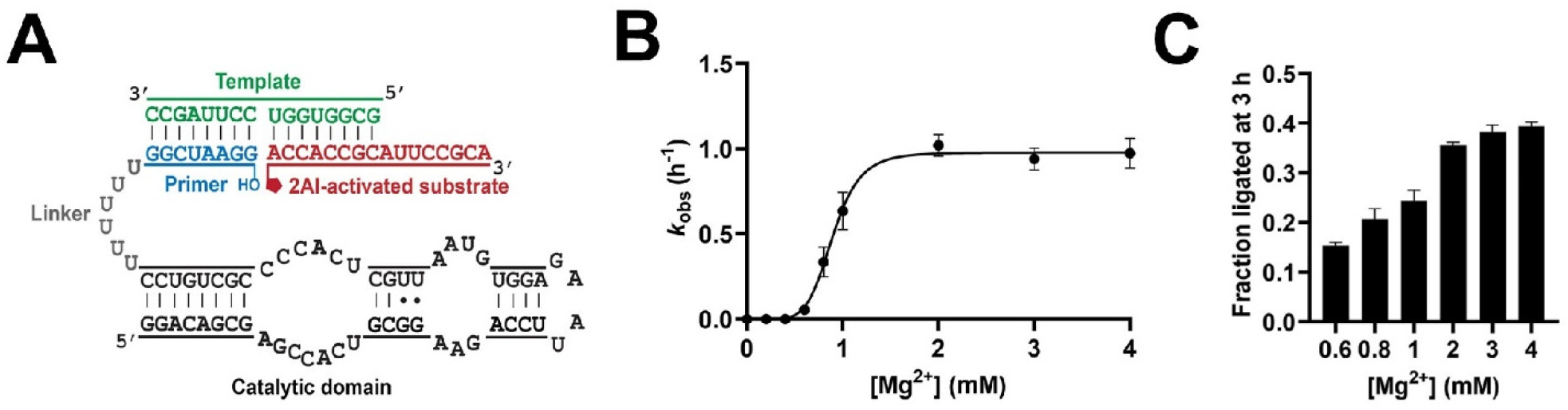
Ligation rates and yields with ribozyme RS8_trunc in response to various Mg^2+^ concentrations. **A**. Schematic of RNA-catalyzed RNA ligation using RS8_trunc and a 2-aminoimidazole activated substrate. **B**. Ligation is undetectable under 0.6 mM. Ligation rate increases sharply between 0.6 mM and 2 mM and remains roughly unchanged above 2 mM Mg^2+^. **C**. Ligation yields after 3 h increase with an increase in Mg^2+^ concentration up to 40% at 2 mM Mg^2+^ and above). Ligation reactions contained 1 µM ribozyme, 1.2 µM RNA template, and 2 µM 2-AI-activated RNA substrate in 100 mM Tris-HCl (pH 8.0), 250 mM NaCl and the indicated concentrations of MgCl_2_ (0-4 mM).

### Effect of Mg^2+^ on vesicle stability and RNA retention

Having identified a ligase ribozyme that functions at low concentrations of Mg^2+^ and in the absence of Na^+^, we set out to test the ability of fatty acid vesicles to retain an encapsulated ribozyme and its substrate in the presence of varying Mg^2+^ concentrations. We used vesicles made of a mixture of decanoic acid (DA, C10:0), decanol (DOH), and glycerol monodecanoate (GMD) because the alcohol and glycerol monoester improve the stability of decanoic acid vesicles. Notably, all three components can be generated under prebiotically relevant conditions (39) and glycerol monoesters, which represent a sub-structure of contemporary phospholipids, can also constitute membranous structures themselves. (38, 40) Since prebiotic processes likely produced a mixture of fatty acids with different chain lengths, where the presence of longer chain fatty acids would have favored vesicle self-assembly, (41) we also tested DA/DOH/GMD vesicles with 10% lauric acid (LA, C12:0) and vesicles made of oleic acid (OA, C18:1). We encapsulated fluorescently labeled RNA oligonucleotides corresponding to either the ligase ribozyme construct (RS8_trunc, 70 nt) that contains an 8 nt primer connected to the 56 nt catalytic domain by a 6 nt polyuridine linker or the 16 nt substrate within DA/DOH/GMD (4:1:1, *v/v/v*), DA/DOH/GMD (4:1:1, *v/v/v*)/10% LA, or OA vesicles (Figure 2A and Supplementary Figure S5A) in 250 mM Tris-Cl (pH 8.0). Vesicles were extruded through 100 nm pore Whatman™ membrane filters, tumbled overnight (∼12 h), and separated from unencapsulated RNA by size-exclusion chromatography (Supplementary Figure S6). (38) We then added various concentrations of Mg^2+^ to the RNA-containing vesicles and monitored RNA leakage over time by size exclusion chromatography. As expected, leakage was insignificant in the absence of Mg^2+^ (ref. (33)) but increased in the presence of Mg^2+^, with only 70% of the ribozyme and 30% of the substrate RNA retained within DA/DOH/GMD vesicles in the presence of 2-3 mM Mg^2+^ after 3 h (Figure 2B and Supplementary Figure S5B). The presence of 10% lauric acid improved RNA retention; 95% ribozyme was retained in 3 mM Mg^2+^ after 3 h (Supplementary Figure S7A), highlighting a potential benefit of the innate heterogeneity of prebiotic fatty acid synthesis. Vesicles composed of oleic acid exhibited 83% ribozyme retention after 3 h at 3 mM Mg^2+^ (Supplementary Figure S7A).

**Figure 2.**
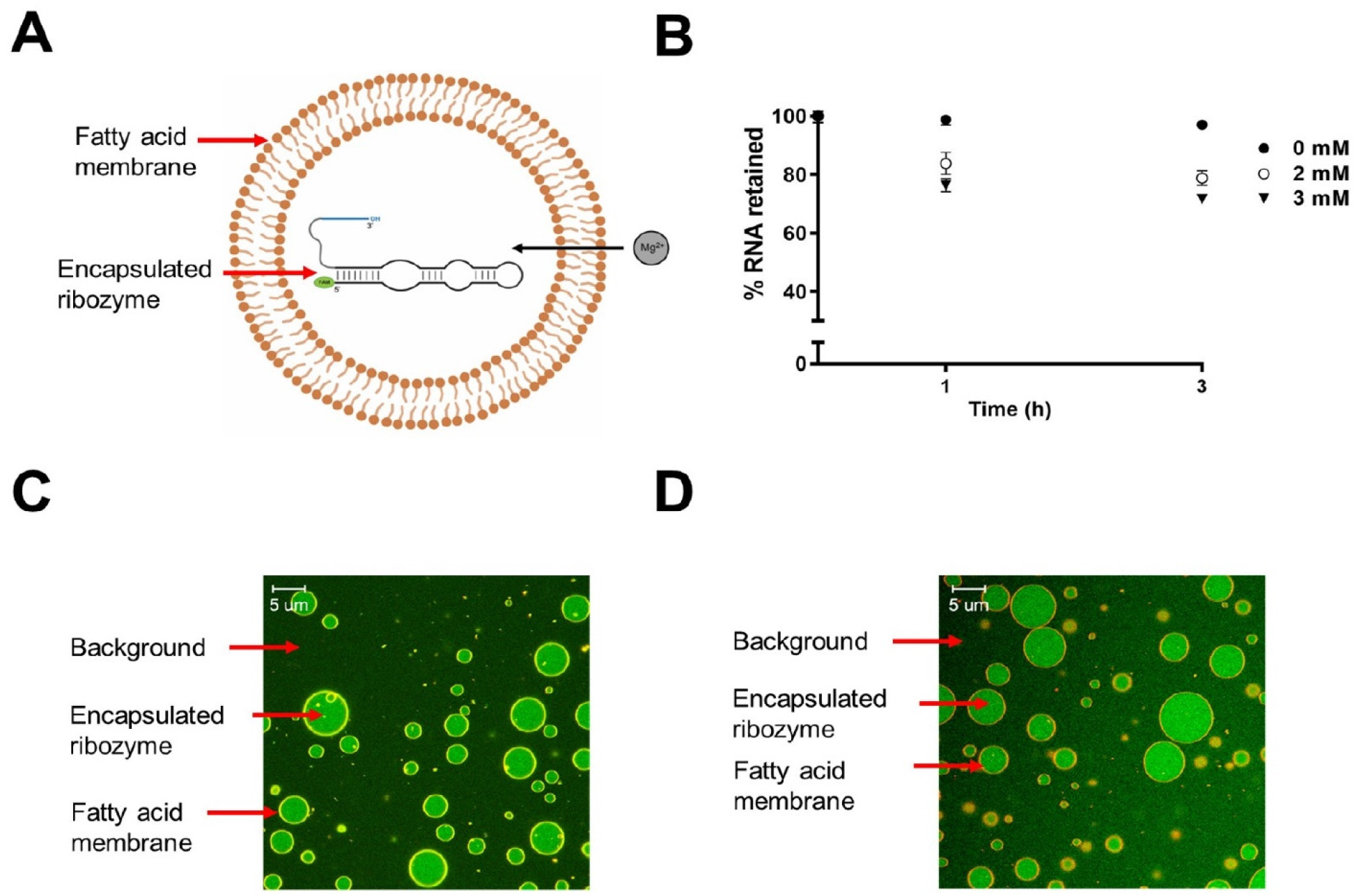
Ribozyme retention within fatty acid vesicles in presence of Mg^2+^. **A**. 5 µM FAM-labeled ligase ribozyme (RS8_trunc; for secondary structure and sequence, see Figure 1A, Supplementary Table S1) was encapsulated within DA/DOH/GMD (4:1:1) vesicles and 2 mM or 3 mM Mg^2+^ was added to its exterior. **B**. Leakage of FAM-labeled ribozyme from vesicles was followed over 3 h. Negligible leakage was observed in the absence of Mg^2+^ and ∼30% leakage was observed in the presence of 2 or 3 mM Mg^2+^ after 3 h. **C, D**. Representative confocal micrographs of DA/DOH/GMD vesicles containing a FAM-labeled ligase ribozyme in the presence of 3 mM Mg^2+^. Membranes were stained using Rhodamine B (red). Vesicles at (C) t = 0 and (D) t = 3 h. Scale bars represent 5 µm.

To enable direct observation of RNA encapsulation and retention by confocal microscopy, we prepared giant unilamellar vesicles (GUVs) made of DA/DOH/GMD with an encapsulated fluorescently labeled ribozyme. In contrast to bulk leakage assays, confocal microscopy allowed us to estimate leakage in individual vesicles and quantify the distribution of leakage in a sample population of vesicles. We diluted the vesicles in 250 mM Tris-Cl (pH 8.0) to dilute the unencapsulated RNA outside of the vesicles. We used fluorescence intensity as a measure of relative concentrations of encapsulated material. To estimate the extent of RNA leakage over time, we compared the fluorescence intensities inside and outside of the vesicles in a representative sample over 3 h (Figure 2C, D and Figure 3) in 3 mM Mg^2+^. At time t = 0, for most vesicles, the ratio of intensities (*I*_*in*_/ *I*_*out*_) ranged from 2.4 to 3.2, with the majority (45%) exhibiting an *I*_*in*_/ *I*_*out*_ value of 2.8 ± 0.1. The maximum decreased with time, indicating gradual leakage of encapsulated RNA. For example, the percentage of vesicles with an *I*_*in*_/ *I*_*out*_ value of 2.8 ± 0.1 decreased to 7% after 1 h, and to zero after 2 h. The peak of the distribution shifted to 2.0 ± 0.1 after 2 h and 1.4 ± 0.1 after 3 h, accompanied by an overall shift in the distribution toward lower *I*_*in*_/ *I*_*out*_ values, indicative of gradual leakage (Figure 3).

**Figure 3.**
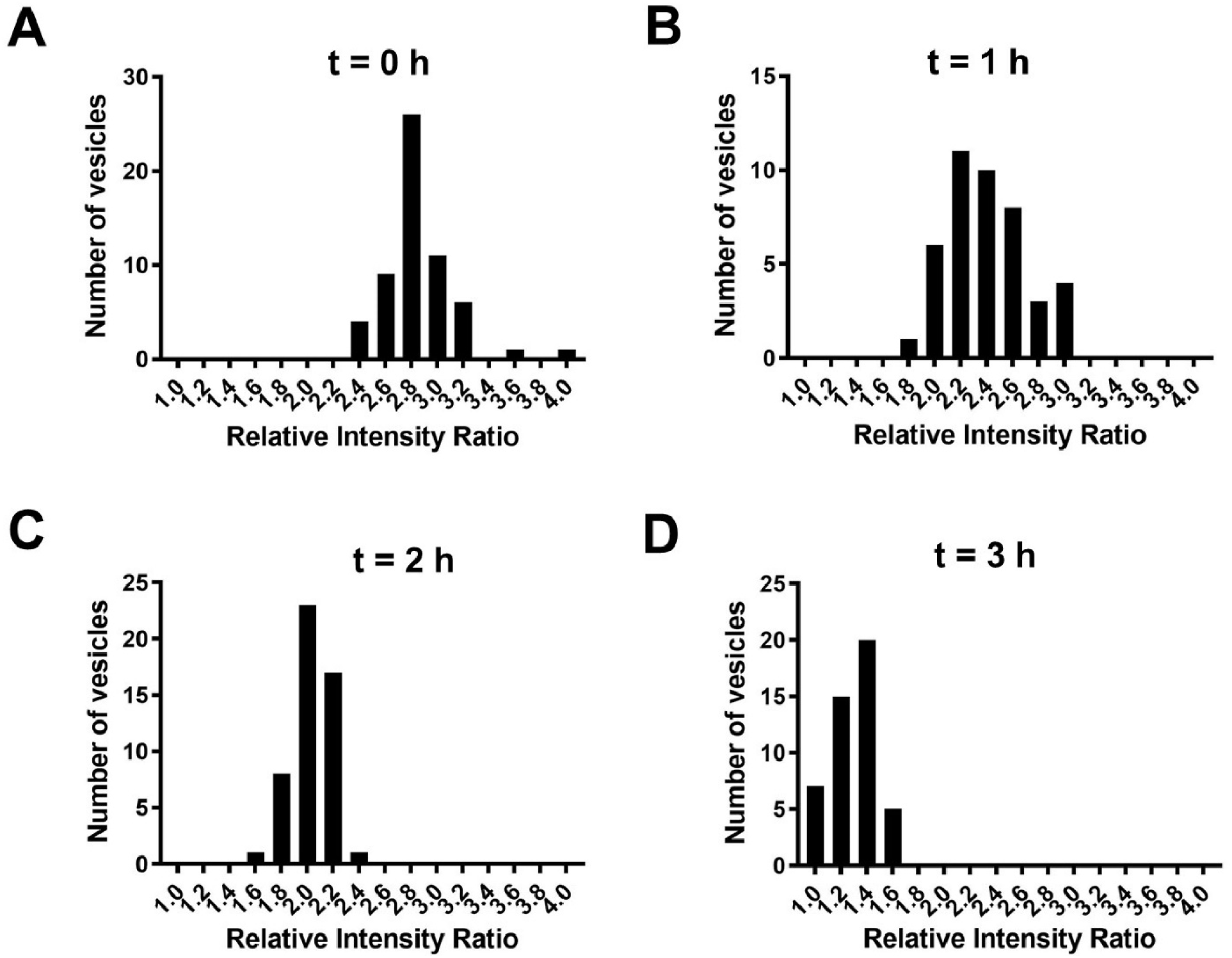
Distribution of RNA leakage in a sample population of DA/DOH/GMD vesicles estimated from confocal micrographs. Confocal micrographs of a population of GUVs were used to monitor ribozyme leakage over 3 h at 3 mM Mg^2+^. At t = 0 (A), the population distribution was centered around an *I*_*in*_/ *I*_*out*_ value of 2.8, where *I*_*in*_/ *I*_*out*_ represents the ratio of the fluorescence intensities inside and outside the vesicle. The distribution shifted left with time (B-D) with most vesicles showing markedly reduced *I*_*in*_/ *I*_*out*_ values, indicating leakage of fluorescently labeled RNA. The distribution at each time point is generated by statistical sampling of GUVs.

Although membrane permeability toward substrates is desirable, retention of ribozymes is essential for protocellular function. The presence of nucleobases and sugars has been shown to decrease flocculation of fatty acid vesicles in the presence of salt. (42) (43) We wondered if association of these small molecules with fatty acid membranes would increase RNA retention in the presence of Mg^2+^. We measured RNA leakage from DA/DOH/GMD vesicles in the presence of D-ribose or adenine. A fluorescently labeled ribozyme was first encapsulated within vesicles. The vesicles were then extruded to 100 nm, after which the vesicles were separated from unencapsulated species by size exclusion chromatography. Next, D-ribose or adenine was added to these purified vesicles and tumbled for 1 h to let the additives interact with the vesicle membrane following which 3 mM Mg^2+^ was added. RNA retention was measured by size exclusion chromatography as discussed above. While the fraction of retained ribozyme fell to ∼70% after 3 h in vesicles without additives, we observed 92% retention in the presence of 0.5% *(w/v*) ribose (∼30 mM) and 89% retention in the presence of 10 mM adenine (Figure 4A). Remarkably, ribose increased substrate retention from 30% to 75% (Figure 4B). Increased RNA retention within primitive cells due to interactions between their membranes and a variety of small molecules present in their local environments could have been a general solution to salt-induced membrane leakage.

**Figure 4.**
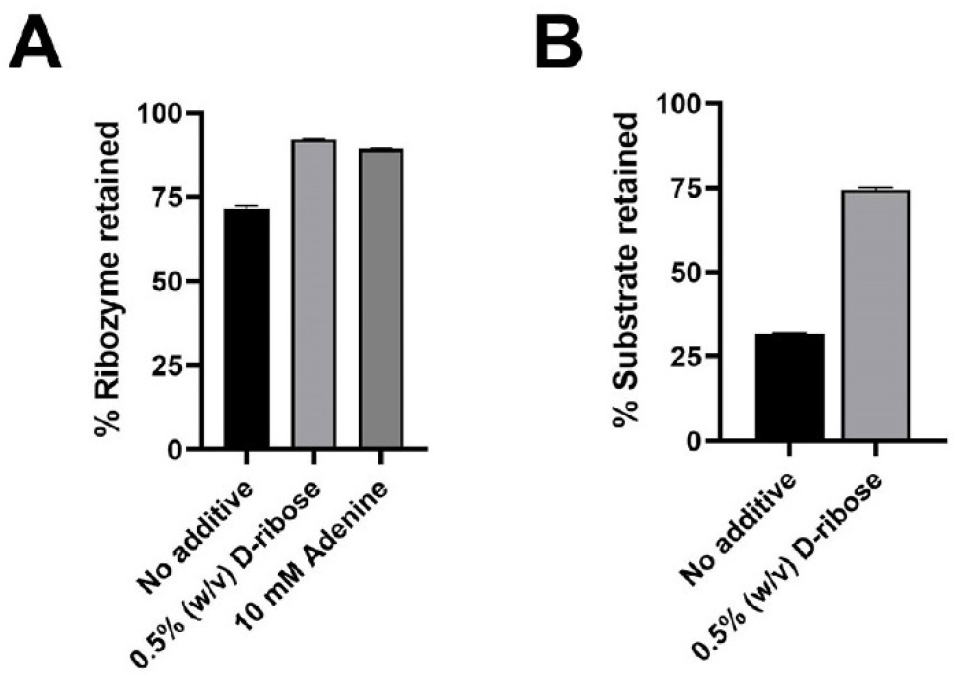
RNA leakage from DA/DOH/GMD vesicles in the presence of D-ribose or adenine obtained from size exclusion chromatography assay. **A**. Leakage of the ribozyme (RS8_trunc) from vesicles composed of DA/DOH/GMD (4:1:1) was monitored over 3 h in the presence of 3 mM Mg^2+^ and D-ribose or adenine. ∼8% leakage was observed in the presence of 0.5% (*w/v*) D-ribose, while the addition of adenine increased leakage to ∼12%. Leakage without additives was ∼30%. **B**. Improved substrate retention was also observed in the presence of 0.5% (*w/v*) D-ribose upon addition of 3 mM Mg^2+^.

### RNA-catalyzed RNA ligation within vesicles

After confirming that the ligase ribozyme and its substrate can be encapsulated and retained within fatty-acid vesicles in the presence of Mg^2+^, we proceeded to examine encapsulated RNA-catalyzed RNA ligation. First, we tested the ligation efficiency of the RS8_trunc ribozyme within DA/DOH/GMD, DA/DOH/GMD/10% LA, and OA vesicles, extruded to 100 nm. We encapsulated RNA sequences representing the fluorescently labeled ribozyme-primer construct (2 μM), template (3 μM), and 2AI-activated substrate (4 μM) (Figure 1A) inside vesicles and purified away the unencapsulated material from the encapsulated fractions using size exclusion chromatography. We added 3 mM Mg^2+^ to the exterior of the vesicles to initiate ligation, collected the encapsulated reactions after 3 h by size exclusion chromatography, and analyzed them by lysing the vesicles and running the RNA on a denaturing PAGE. We observed 18.6%, 10%, and 6.8% ligation within DA/DOH/GMD, DA/DOH/GMD/10% LA, and OA vesicles, respectively (Supplementary Figure S7B). Although the reason for reduced ligation yields within DA/DOH/GMD/10% LA, and OA vesicles is not clear, we suggest that it could be a result of chelation of Mg^2+^ by these lipids. (44) Adding 150 mM free fatty acids to an unencapsulated ribozyme reaction mixture eliminated ligation in the presence of 3 mM Mg^2+^ (Supplementary Figure S7B), consistent with chelation of Mg^2+^ by free fatty acids. (45)

As ligation yield was the highest within DA/DOH/GMD vesicles, we selected these vesicles to study the kinetics of compartmentalized RNA-catalyzed RNA ligation. We encapsulated the ribozyme, template, and substrate, extruded and purified vesicles containing encapsulated RNA, and initiated catalysis by adding 2 mM or 3 mM Mg^2+^ to the vesicles. Reaction aliquots were analyzed at various time points over 3 h. We observed a steady increase in ligation over time with 17.5% and 18.6% ligation after 3 h in 2 mM and 3 mM Mg^2+^, respectively (Figure 5A, C). This was comparable to ligation yields outside vesicles (i.e., in solution) which were 21.6% and 26% in 2 mM and 3 mM Mg^2+^, respectively (Figure 5B, C) (surprisingly, the ligation efficiency of FAM labeled RS8_trunc was lower than that of its unlabeled counterpart in solution (Supplementary Figure S8)). *k*_obs_ values for encapsulated ligation were 0.72 h^-1^ and 0.79 h^-1^ in 2 mM and 3 mM Mg^2+^, respectively whereas *k*_obs_ values for ligation in solution were 0.63 h^-1^ and 0.64 h^-1^ in 2 mM and 3 mM Mg^2+^, respectively (Figure 5A, B, D, E).

**Figure 5.**
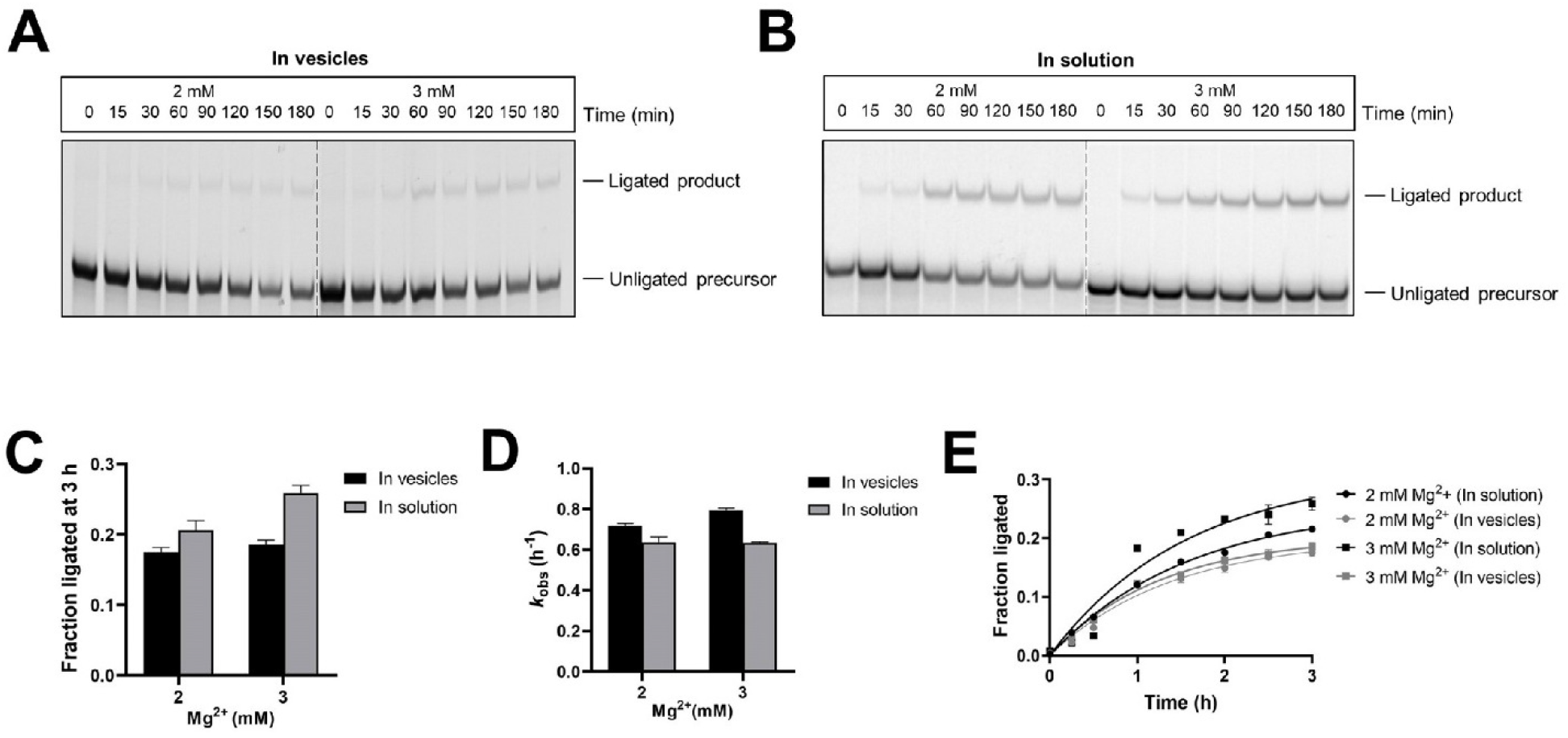
RNA-catalyzed RNA ligation within fatty acid vesicles. **A**. RNA-catalyzed ligation inside DA/DOH/GMD vesicles at 2 and 3 mM Mg^2+^. **B**. RNA-catalyzed ligation in solution at 2 and 3 mM Mg^2+^. **C**. Ligation yields at 3 h in vesicles were comparable to the yields in solution. **D**. Ligation rates within vesicles were comparable to the rates in solution at 2 mM and 3 mM Mg^2+^, respectively. **E**. Ligation kinetics in 2 mM and 3 mM Mg^2+^ in solution and when encapsulated within DA/DOH/GMD vesicles. Ligation reactions containing 2 µM FAM-labeled ribozyme (RS8_trunc), 3 µM RNA template, 4 µM 2AI-activated substrate, and 100 mM Tris-HCl (pH 8.0) were encapsulated within vesicles, the encapsulated fractions were purified away from the unencapsulated fractions, and 2 mM or 3 mM Mg^2+^ was added to the encapsulated fraction to initiate the reactions. Aliquots at different time points were re-purified by size-exclusion chromatography and the encapsulated RNA was analyzed by denaturing PAGE.

As addition of ribose led to a marked reduction of RNA leakage (Figure. 4), we repeated the encapsulated ligation reaction in the presence of 0.5% (*w/v*) ribose at 3 mM Mg^2+^ by first adding ribose to the RNA-containing vesicles, tumbling for 1 h, and then adding 3 mM Mg^2+^. While ligation yield increased from 18% to 28%, to our surprise *k*_obs_ fell ∼2-fold to 0.38 h^-1^ in ribose-containing vesicles. We observed a similar trend for the corresponding unencapsulated ligation reaction in the presence of ribose, where ribose caused an increase in ligation yield from 26% to 38%, but a reduction in *k*_obs_ from 0.64 h^-1^ to 0.49 h^-1^ (Figure 6). The increase in ligation yield in both the encapsulated and unencapsulated reactions in the presence of ribose could be due to an increase in the fraction of correctly folded ribozymes as a result of ribose-induced molecular crowding effects. (46) However, ribozymes folded in presence of ribose exhibit reduced rates, for reasons that are unclear. Nonetheless, these effects are modest, and the extent of RNA-catalyzed RNA ligation, both in the presence and absence of ribose, was comparable inside and outside vesicles (Figure 5 and Figure 6).

**Figure 6.**
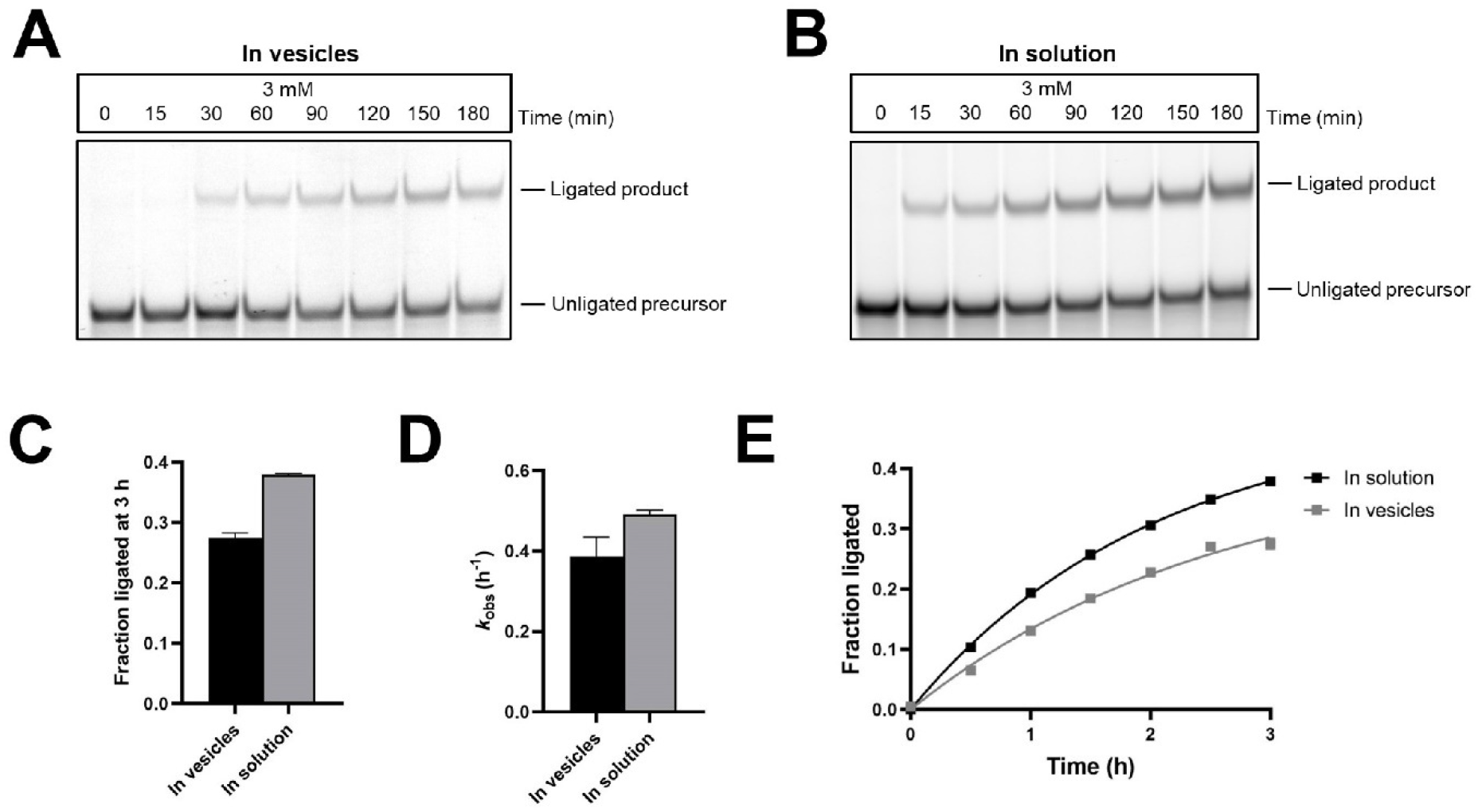
RNA-catalyzed RNA ligation within fatty acid vesicles in the presence of D-ribose. **A**. RNA-catalyzed ligation inside DA/DOH/GMD vesicles in the presence of 0.5% (*w/v*) D-ribose at 3 mM Mg^2+^. **B**. RNA-catalyzed ligation in solution in the presence of 0.5% (*w/v*) D-ribose at 3 mM Mg^2+^. **C**. Ligation yields after 3 h in vesicles were comparable to yields in solution at 3 mM Mg^2+^ in the presence of ribose. **D**. Ligation rates within vesicles were comparable to the rates in solution at 3 mM Mg^2+^ in the presence of ribose. **E**. Ligation kinetics in the presence of 0.5% (*w/v*) D-ribose in solution and when encapsulated within DA/DOH/GMD vesicles. Ligation reactions containing 2 µM FAM-labeled ribozyme (RS8_trunc), 3 µM RNA template, 4 µM 2AI-activated substrate, and 100 mM Tris-HCl (pH 8.0) were encapsulated within vesicles, and the encapsulated fractions were purified away from the unencapsulated fractions. The encapsulated fractions were tumbled with 0.5% (*w/v*) D-ribose for 1 h and 3 mM Mg^2+^ was added to the to initiate the reactions. Aliquots at different time points were re-purified by size-exclusion chromatography and the encapsulated RNA was analyzed by denaturing PAGE.

## Conclusion

Compartmentalized RNA catalysis is fundamental to our conception of the emergence and evolution of RNA-based early life. RNA-catalyzed RNA assembly within protocells is thought to have been especially important for efficient genome replication and the emergence of catalytic molecules in primordial biology. The primary roadblock toward constituting ribozyme-catalyzed reactions, including RNA assembly, within prebiotic fatty acid compartments has been the incompatibility between the Mg^2+^ requirement for ribozyme function and the instability of short chain fatty acid vesicles in the presence of low to moderate concentrations of Mg^2+^. High concentrations of Mg^2+^ also catalyze RNA hydrolysis and the hydrolysis of labile phosphorimidazolides. The tendency of fatty acid vesicles to leak in salty environments has led to recent speculations about the relevance of more Mg^2+^-resistant cyclophospholipid vesicles to early life. (47, 48) However, the diffusion of nutrients such as RNA building blocks across cyclophospholipid membranes and the growth and division of protocells bounded by cyclophospholipids are yet to be demonstrated.

In this work, we provide a partial resolution to the apparent conflict between RNA catalysis and protocell stability by identifying a ligase ribozyme that functions at low Mg^2+^ concentrations and demonstrating that prebiotic small molecules such as ribose and adenine reduce RNA leakage from protocells in the presence of Mg^2+^. These discoveries allowed us to constitute ribozyme-catalyzed RNA ligation of a 2AI-activated RNA substrate inside prebiotically relevant DA/DOH/GMD vesicles. We found that ligation yields and rates are comparable (within two-fold) both inside and outside vesicles in the presence of 2 mM or 3 mM Mg^2+^, suggesting that fatty acid protocells may support ribozyme-catalyzed RNA assembly, while keeping the products of the reaction separated from the surrounding environment.

The greater permeability of fatty acid membranes to the 16 nt RNA substrate compared to the 70 nt ribozyme-primer construct presents an attractive model for compartmentalized RNA-catalyzed RNA copying. In this model, short oligomeric substrates are generated by nonenzymatic polymerization/ligation of 2AI-activated nucleotides outside protocells. (49-51) These oligonucleotides could then be internalized through temporary defects in the protocell membrane induced by Mg^2+^, and utilized in copying longer RNA templates by ribozymes. Since longer RNAs are retained inside protocells, the beneficial effects of functional RNAs generated by RNA-catalyzed assembly within protocells would be restricted to the same protocell, and the accumulation of these beneficial effects would eventually result in a population of spontaneously evolving protocells. Despite the widely recognized importance of ribozyme function within fatty acid protocells in modeling early cellular life, there has been just a single example in the form of endoribonucleolytic cleavage by the hammerhead ribozyme inside prebiotically implausible myristoleic acid vesicles. (38, 52) Our demonstration of compartmentalized RNA-catalyzed RNA ligation provides the first instance of RNA assembly catalyzed by ribozymes within prebiotically plausible fatty acid-bound model protocells.

More broadly, the importance of ribozyme function under low [Mg^2+^] conditions is highlighted by the ribozymes found in biology, which have evolved to function under cellular conditions that provide only 0.5-1 mM free Mg^2+^. However, it is not clear how abundant ribozymes that function in the low Mg^2+^ regime are in the RNA sequence space. In the absence of sufficient representation of such ribozymes in the RNA world, more general phenomena such as molecular crowding may have been crucial for lowering the Mg^2+^ requirements of ribozymes. (46) The first ribozymes must have emerged from nonenzymatic RNA assembly processes, which to our knowledge, require high Mg^2+^ concentrations. To accommodate compartmentalized nonenzymatic RNA assembly, early protocell membranes may have been composed of heterogeneous mixtures of lipids that are more resistant to Mg^2+^-induced destabilization. After the emergence of robust RNA-catalyzed RNA assembly with ribozymes that are active at low Mg^2+^, compartmentalized RNA assembly could spread to more diverse environments with low Mg^2+^ concentrations, such as freshwater ponds, where simpler and more readily available fatty acids might have sufficed for cell membrane assembly. (53)

Over evolutionary time, single chain fatty acid vesicles would have gradually transitioned to diacylphospholipid vesicles, which are chemically closer to modern cells. While diacylphospholipid vesicles are stable in the presence of high Mg^2+^ concentrations, (54) they are impermeable to RNA monomers and even Mg^2+^. (33, 54) Protocells with heterogenous membranes composed of both fatty acids and phospholipids, which retain moderate permeability while imparting stability in challenging ionic environments, could be an intermediate stage in protocellular evolution. (54) Until the emergence of such chemically complex salt resistant protocell membranes, ribozymes with low Mg^2+^ requirements could have been important in driving genome replication and protocellular metabolism.

## Experimental Section

Materials and methods are described in detail in the Supporting Information.

## Acknowledgment

J.W.S. is an Investigator of the Howard Hughes Medical Institute. This work was supported in part by a grant from the Simons Foundation (290363) to J.W.S. The authors thank Dr. Harry Aitken, Dr. Longfei Wu, Dr. Lijun Zhou, and Xiwen Jia for helpful comments on the manuscript.

## Table of Contents Graphic

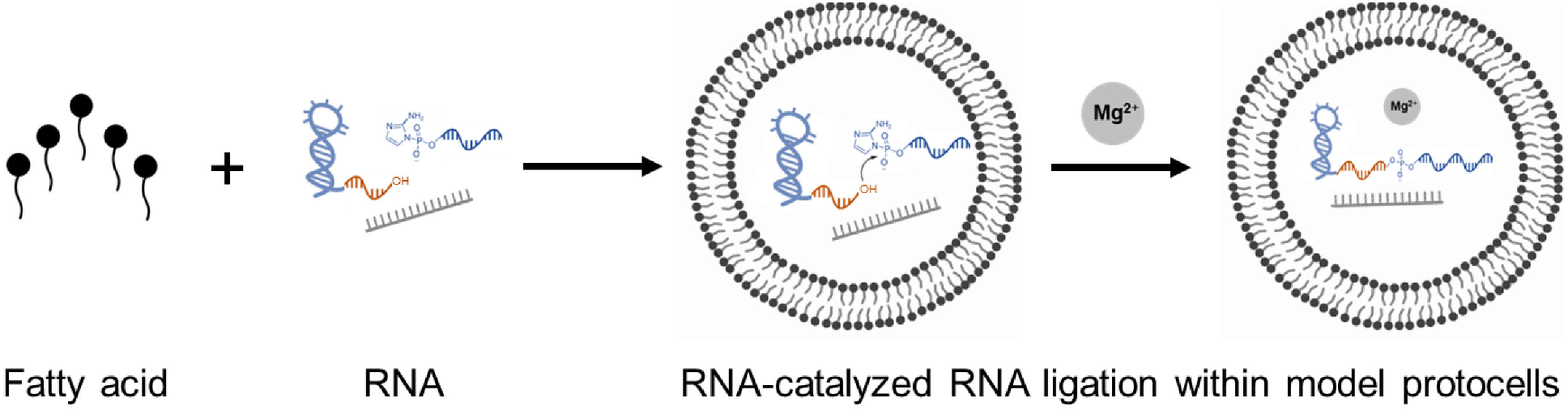

