## Supporting Information for "RNA-catalyzed RNA Ligation within Prebiotically Plausible Model Protocells"

**Table of Contents:**

Experimental Methods……………………………………………………………………….3

Supplementary Table S1…………………………………………………………………….7

Supplementary Figures S1 to S8……………………………………………………..….8

References…………………………………………………………………………..…..….16

Author Contributions…………………………………………………………..………16

### Experimental Methods

#### Materials

Single stranded DNA used to generate dsDNA templates for transcription, 5′-monophosphate RNAs, and fluorescently labeled RNAs were purchased from Integrated DNA Technologies (IDT). Decanoic acid (DA, C10:0), 1-decanol (DOH), monocaprin (the glycerol monoester of decanoic acid, GMD) and oleic acid (OA, 18:1) were purchased from Nu-Chek Prep. Lauric acid (LA, C12:0) was purchased from ACROS Organics. Rhodamine B and Sepharose 4B were purchased from Sigma-Aldrich. 10 mL disposable chromatography columns were purchased from BioRad Laboratories. D- (–)-ribose and adenine were purchased from Sigma-Aldrich.

#### Ribozyme preparation

Unlabeled ligase ribozymes were prepared by *in vitro* transcription of dsDNA templates generated by PCR. The template strand contained 2′-*O*-methyl modifications on the last two nucleotides to reduce transcriptional heterogeneity at the 3′-end of the RNA[1] (Table S1). Transcription reactions were carried out in the presence of 40 mM Tris-HCl (pH 8), 2 mM spermidine, 10 mM NaCl, 25 mM $MgCl_2$, 10 mM dithiothreitol (DTT), 30 U/mL RNase inhibitor murine (NEB), 2.5 U/mL thermostable inorganic pyrophosphatase (TIPPase) (NEB), 4 mM of each NTP, 30 pmol/mL DNA template, and 1 U/µL T7 RNA Polymerase (NEB) for 3 h at 37°C. The reaction was quenched with DNase I (NEB) and RNA was extracted with phenol-chloroform-isoamyl alcohol (PCI), ethanol precipitated, and purified by denaturing PAGE.

#### 2-aminoimidazole activation of the 5′-phosphorylated RNA substrate

The 2-aminoimidazole-activated RNA substrate was generated from the corresponding 5′-monophosphorylated oligonucleotide by incubating it with 0.2 M 1-ethyl-3-(3 dimethylaminopropyl) carbodiimide (HCl salt) and 0.6 M 2-aminoimidazole (HCl salt, pH adjusted to 6) for 2 h at room temperature. The product was washed with water in Amicon Ultra spin columns (3 kDa cutoff) five times (200 µL milli-Q water per wash) and purified by reverse phase analytical HPLC using a gradient of 98% to 75% 20 mM TEAB (triethylamine bicarbonate, pH 8) versus acetonitrile over 40 min.[2]

#### Ligation reactions outside vesicles

Ligation reactions (5 µL) performed outside vesicles contained 1 µM ribozyme, 1.2 µM RNA template, and 2 µM 2-AI-activated RNA substrate in 100 mM Tris-HCl (pH 8.0), with various concentrations of NaCl (0-250 mM) and $MgCl_2$ (0-5 mM). 1 µL aliquots taken at various times were quenched with 5 µL quench buffer (8M urea, 100 mM Tris-Cl, 100 mM boric acid, 100 mM EDTA) and analyzed by denaturing PAGE. Gels were stained using SYBR™ Gold, imaged on a Typhoon 9410 scanner (GE Healthcare, Little Chalfont, Buckinghamshire, UK), and analyzed in ImageQuant IQTL 8.1. Kinetic plots were nonlinearly fitted (using GraphPad Prism 8.4.0) to the modified first order rate equation, y = A (1 – $e^{-kx}$), where A represents the fraction of active complex, *k* is the first order rate constant, x is time, and y is the fraction of ligated product.

**Preparation of giant unilamellar vesicles (GUVs) for microscopy**
200 mM fatty acid micelles were prepared by dissolving 100 µmol of neat mixed amphiphiles composed of DA:DOH:GMD (4:1:1, v/v/v) in 1.25 equivalent of NaOH solution to a volume of 500 µL with Milli-Q water. Fatty acid GUVs were prepared by resuspending the micelle solution in buffer stock (1 M PIPES, pH ~6.6) with fluorescently labeled RNA, and water to a final concentration of 50 mM decanoic acid, and 100 mM PIPES buffer as described previously.[3] The membrane was stained with 10 µM rhodamine B (red). Confocal images were collected using a Nikon A1R HD25 confocal laser scanning microscope equipped with LU-N4/N4S 4-laser unit and visualized in Image J (FIJI).

**RNA encapsulation within small unilamellar vesicles (SUVs) for bulk leakage assays**
Small unilamellar vesicles were prepared by mixing DA, DOH, and GMD 4:1:1, v/v/v) at a total lipid concentration of 150 mM with buffer (250 mM Tris-HCl, pH 8.0) to a total volume of 250 µL as described previously.[4] Encapsulation of fluorescently labeled RNA was achieved by mixing RNA with buffer before addition to the neat amphiphiles. Vesicles were extruded through 100 nm pore Whatman™ membrane filters using a MiniExtruder system (Avanti Polar Lipids), and then allowed to tumble overnight (>12 h).

**Purification of SUVs containing encapsulated RNA**
*Preparation of size exclusion chromatography columns.* 5 mL of an ethanolic Sepharose 4B slurry was added to a disposable 10 mL chromatography column and allowed to settle until ethanol approached the top of the resin bed. Ethanol was removed by applying 5 mL of deionized water to the top of the resin three to five times and allowing it to flow through the column. 5 mL of 250 mM Tris-HCl (pH 8.0) was then added to the top of the resin and allowed to flow through two times before use. The running buffer containing 50 mM DA/DOH/GMD or DA/DOH/GMD/10% LA, or 5 mM OA with a composition matching the membranes of the purified vesicles was used to ensure that the total fatty acid concentration remained above the critical aggregation concentration of the amphiphile, which is necessary to prevent disruption of vesicles during purification.[4, 5] The running buffer was filtered through a 0.22 µm syringe filter unit before use, to remove any potential aggregates. The running buffer was then kept at 1 ml above the top of the resin until the sample was ready to be added.

*Collection of size exclusion chromatography fractions.* The extruded SUVs were gradually loaded to the resin bed without disturbing the resin surface. After the SUVs were well into the resin, running buffer was added dropwise until the unencapsulated solute layer had reached the middle of the resin. This was followed by the addition of 5 mL of 250 mM Tris-HCl (pH 8.0). The eluent was fractionated and collected using a fraction collector (Gilson FC203B) on a 96-well plate (Corning Costar #3615). The 96-well plate was then read by a microplate reader (SpectraMax i3) using the corresponding excitation and emission wavelengths based on the fluorescent dye attached to the encapsulated RNA.

Note that vesicles elute first, followed by the unencapsulated fraction. The vesicle fraction was collected and tumbled for 16 h before use.

**Quantification of RNA leakage from vesicles**
Extruded vesicles with encapsulated RNA were prepared with 5 µM fluorescently labeled RNA oligonucleotides and purified to remove unencapsulated RNA as described above. To assay leakage in the presence of additives, indicated concentrations of D(–)-ribose or adenine were added to the purified RNA-containing vesicles and tumbled at room temperature for 1 h. $MgCl_2$ was added to the vesicles to initiate leakage. Extent of leakage was measured by size exclusion chromatography, with fluorescence in the encapsulated and non-encapsulated fractions measured with a microplate plate reader. The reported fraction encapsulated over time represents the fraction of the fluorescence remaining in the vesicle fraction after column purification.

To estimate RNA leakage by confocal microscopy, following addition of 3 mM $Mg^{2+}$, a 5 µL vesicle sample was pipetted onto a glass microscope slide, with a cover slip placed on top and secured by clear vacuum grease. At each time point, the sample was imaged by confocal microscopy. Each confocal micrograph was analyzed by ImageJ (FIJI), and the relative fluorescence intensity of each vesicle lumen was compared to the external medium. The resulting distribution of intensity ratios was plotted as a histogram using GraphPad Prism 8.4.0.

**Ligation reactions inside vesicles**
Ligation reactions (100 µL) contained 2 µM FAM-labeled ribozyme (RS8_trunc) (Figure 1, Supplementary Table S1), 3 µM template, 4 µM 2AI-activated substrate, and 100 mM Tris-HCl pH 8.0. The amphiphilic mixture composed of fatty acid, glycerol monoester, and alcohol at a total concentration of 150 mM at the desired component ratio was prepared and added to the ligation reaction mixture before high-speed (>3000 r.p.m.) vortexing for 4-5 s. Vesicles were extruded through 100 nm pore Whatman$^{TM}$ membrane filters using a MiniExtruder system (Avanti Polar Lipids), and then allowed to tumble overnight (>12 h). The vesicle fraction containing encapsulated RNA was purified away from unencapsulated solutes by size exclusion chromatography and collected as described above. For encapsulated ligation reactions in the presence of ribose, 0.5% (*w/v*) D(–)-ribose was added to the purified RNA-containing vesicles and tumbled at room temperature for 1 h. Ligation was initiated by adding $MgCl_2$ to vesicles (with or without ribose addition) and the samples were tumbled at room temperature throughout the course of the reaction. At the indicated time points, aliquots of vesicles (50 µL) were removed and re-purified by size-exclusion chromatography, with 50 mM filtered lipid with matching membrane composition as the mobile phase. The vesicle fractions were collected in a 1.5 mL Eppendorf tube to which Triton (0.1% v/v), glycogen (to a final concentration of 6 µg/ml), and 0.5 mL of cold ethanol was added. The sample was incubated at -20 °C for 2 h, centrifuged at 16.1×1000 r.c.f for 30 min, and supernatant was removed. The pellet was washed with 70% cold ethanol and centrifuged again for 25 min. The supernatant was removed, and the pellet was suspended in 50 µL quench buffer (8 M urea, 100 mM Tris-Cl, 100 mM boric acid, 100 mM EDTA) and analyzed by denaturing PAGE. Gels were imaged on a Typhoon 9410 scanner (GE Healthcare, Little

Chalfont, Buckinghamshire, UK), and analyzed in ImageQuant IQTL 8.1. Kinetic plots were nonlinearly fitted (using GraphPad Prism 8.4.0) to the modified first order rate equation, $y = A\,(1 - e^{-kx})$, where A represents the fraction of active complex, $k$ is the first order rate constant, x is time, and y is the fraction of ligated product.

**Supplementary Table**

| Name | Sequence (5′ → 3′) |
|---|---|
| RS1 | GACUCACUGACACAGAUCCACUCACGGACAGCGGAAUGCUGCCA ACCGUGCGGGCUAAUUGGCAGACUGAGCUCGCUGUCCUUUUUU GGCUAAG**G** |
| RS5 | GACUCACUGACACAGAUCCACUCACGGACAGCGGAACCCUUAUC ACAGUCGUGCGGAUUUGUAAGCCUAAGCGCGCUGUCCUUUUUU GGCUAAG**G** |
| RS8 | GACUCACUGACACAGAUCCACUCACGGACAGCGAGCCACUGCGG AAGACCUUAAGAGGUGUAAUUGCUCACCCCGCUGUCCUUUUUU GGCUAAG**G** |
| RS8_trunc | GGACAGCGAGCCACUGCGGAAGACCUUAAGAGGUGUAAUUGCUC ACCCCGCUGUCCUUUUUUGGCUAAG**G** |
| 2AI-activated substrate | (5′-phosphoro-2AI)-ACCACCGCAUUCCGCA |
| Template | GCGGUGGUCCUUAGCC |

**Table S1. RNA oligonucleotides used in this study.** Fixed base paired stem nucleotides are underlined. Primer sequence is shown in blue, polyuridine linker is shown in gray, and the nucleotide containing the 3′-hydroxyl nucleophile is boldfaced.

### Supplementary Figures

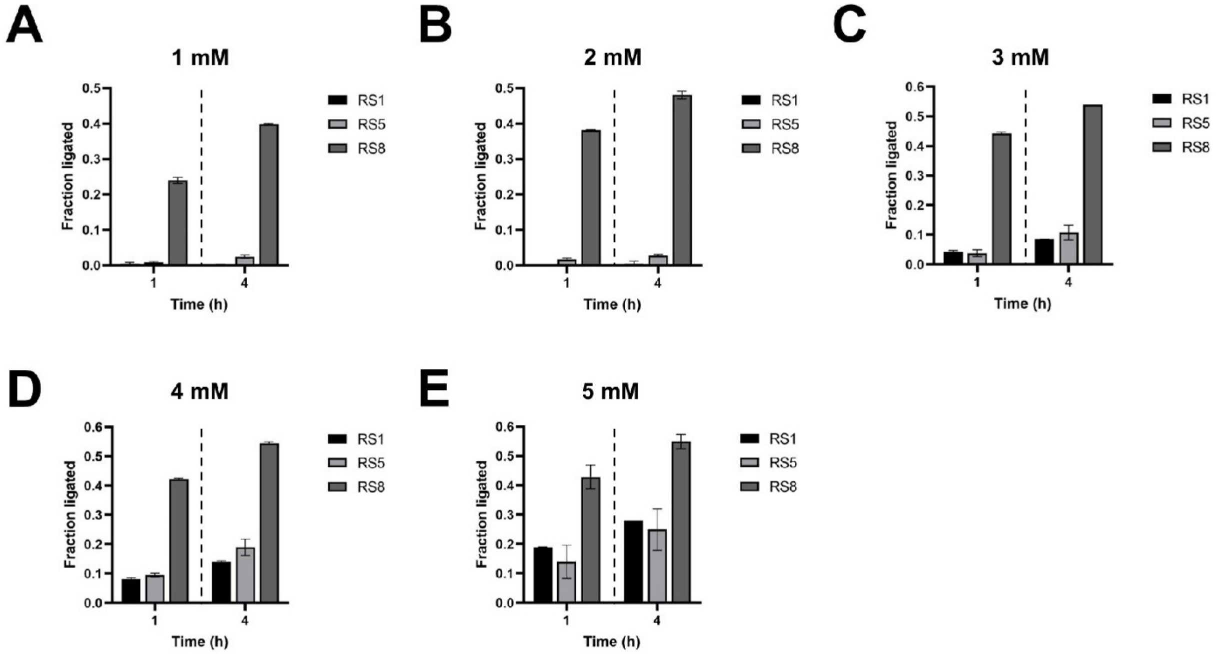


**Figure S1. Screening for ligase ribozyme activity at 1-5 mM $Mg^{2+}$.** Ligation yields with ribozymes, RS1, RS5, and RS8 in the presence of (A) 1 mM $Mg^{2+}$ (B) 2 mM $Mg^{2+}$ (C) 3 mM $Mg^{2+}$ (D) 4 mM $Mg^{2+}$ (E) 5 mM $Mg^{2+}$ after 1 h and 4 h. (See Fig. S2 for representative gels). Ligation reactions contained 1 µM ribozyme, 1.2 µM RNA template, and 2 µM 2-AI-activated RNA substrate in 100 mM Tris-HCl (pH 8.0), 250 mM NaCl, and the indicated concentrations of $MgCl_2$ (0-5 mM).

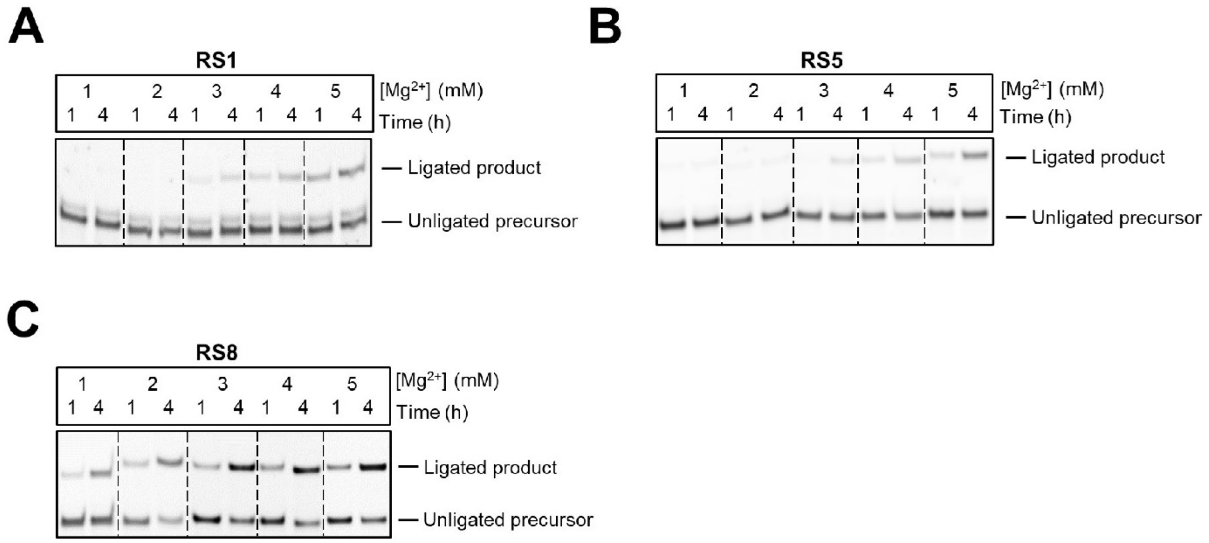


**Figure S2. RNA-catalyzed RNA ligation in the presence of 1-5 mM $Mg^{2+}$. A.** RS1 **B.** RS5 **C.** RS8. Only RS8 has detectable activity at all $Mg^{2+}$ concentrations between 1-5 mM. Ligation reactions contained 1 µM ribozyme, 1.2 µM RNA template, and 2 µM 2-AI-activated RNA substrate in 100 mM Tris-HCl (pH 8.0), 250 mM NaCl, and the indicated concentrations of $MgCl_2$ (0-5 mM).

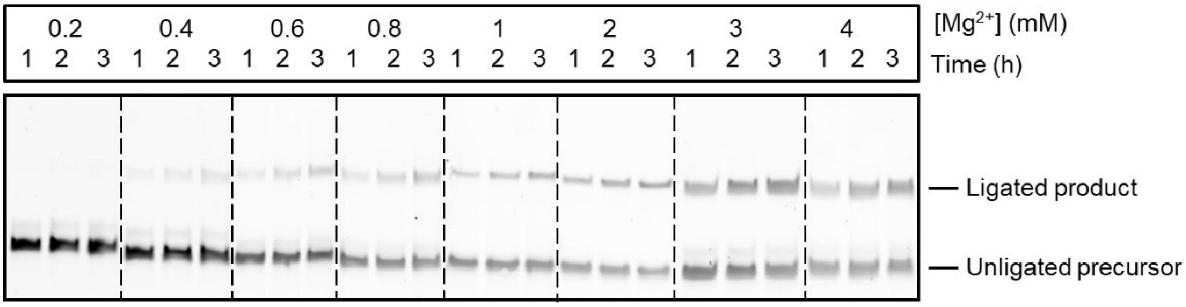


**Figure S3. RNA ligation catalyzed by RS8_trunc in the presence of 0.2-4 mM $Mg^{2+}$.** Significant ligation was detected at 0.6 mM $Mg^{2+}$, which plateaus at 2 mM $Mg^{2+}$. Ligation reactions contained 1 µM ribozyme, 1.2 µM RNA template, and 2 µM 2-AI-activated RNA substrate in 100 mM Tris-HCl (pH 8.0), 250 mM NaCl, and the indicated concentrations of $MgCl_2$ (0-4 mM).

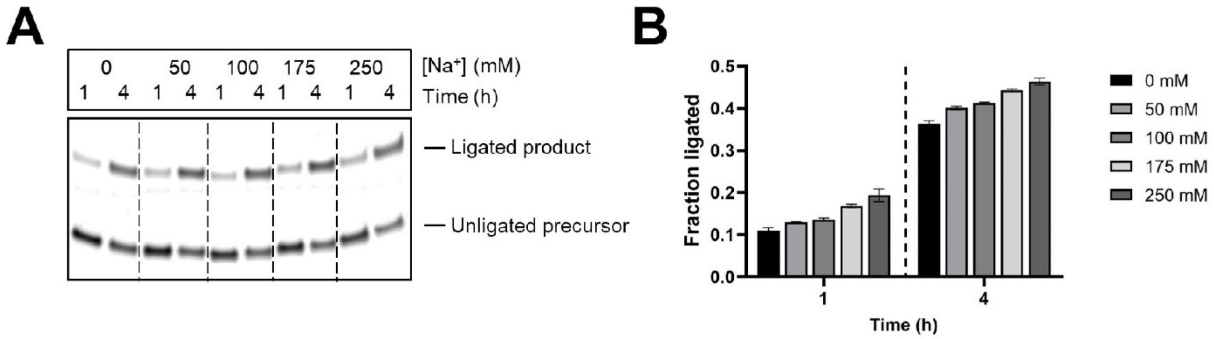


**Figure S4. RNA ligation catalyzed by RS8_trunc in the presence of 0-250 mM $Na^+$.** **A.** Significant ligation was detected in the absence of $Na^+$. **B.** Ligation yields were comparable in the concentration range of $Na^+$ (0-250 mM) tested. Ligation reactions contained 1 µM ribozyme, 1.2 µM RNA template, and 2 µM 2-AI-activated RNA substrate in 100 mM Tris-HCl (pH 8.0), 3 mM $MgCl_2$ and the indicated concentrations of NaCl (0-250 mM).

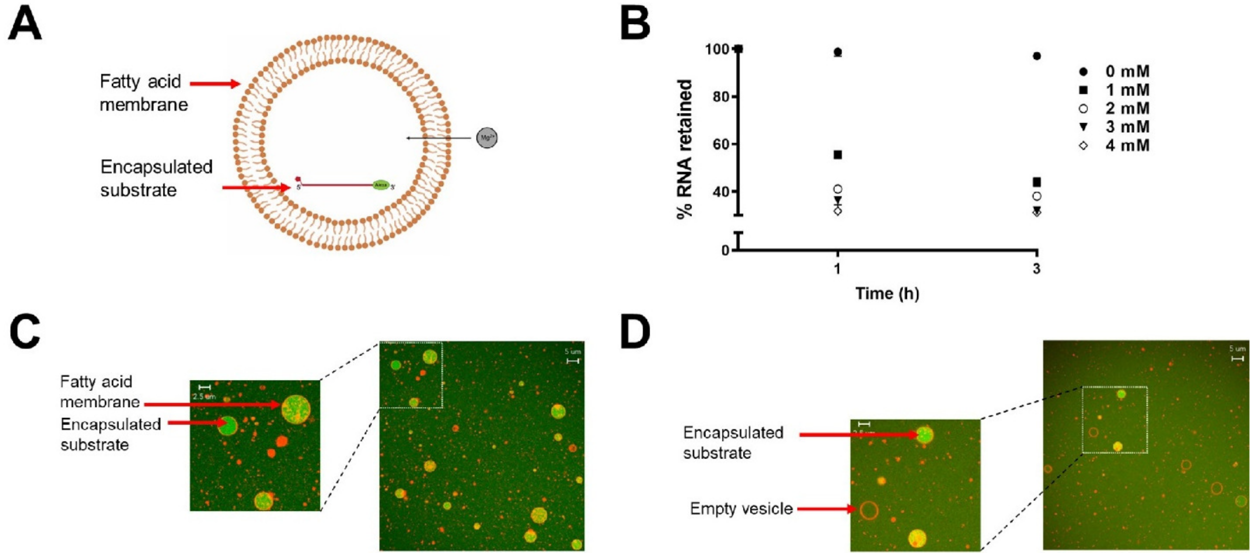


**Figure S5. Stability of fatty acid vesicles and substrate retention in the presence of $Mg^{2+}$. A.** 5 µM Alexa488-labeled RNA oligomer (16-nucleotide-long, green) corresponding to the ligase substrate (Table S1) was encapsulated within DA/DOH/GMD (4:1:1) vesicles and the indicated concentrations of $Mg^{2+}$ was added to its exterior. **B.** Leakage of Alexa488-labeled substrate from vesicles was followed over 3 h. Negligible leakage was observed in the absence of $Mg^{2+}$; however, >70% RNA was lost after 3 h in the presence of 1-3 mM $Mg^{2+}$. **C, D.** Representative confocal micrographs of DA/DOH/GMD vesicles containing the Alexa488-labeled RNA oligomer in the presence of 3 mM $Mg^{2+}$. Membranes were stained using Rhodamine B (red). Vesicles at (C) t = 0 and (D) t = 3 h. Scale bars represent 2.5 um (left) and 5 µm (right).

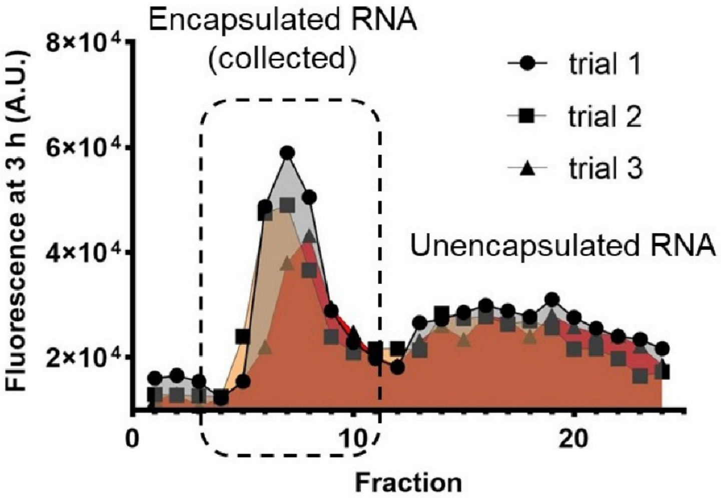


**Figure S6. Representative size exclusion chromatography trace showing the separation of vesicles containing RNA from unencapsulated RNA in solution.** Encapsulated RNA oligonucleotides were purified away from free RNA in solution using sepharose 4B size exclusion chromatography. Vesicles encapsulating RNA, collected in the first fraction, were used for leakage and ligation studies. Leakage was calculated by quantifying fluorescence in the vesicle and free RNA fractions. The first sharp peak in the fluorescence trace represents vesicles containing encapsulated RNA and the second broad peak represents unencapsulated RNA.

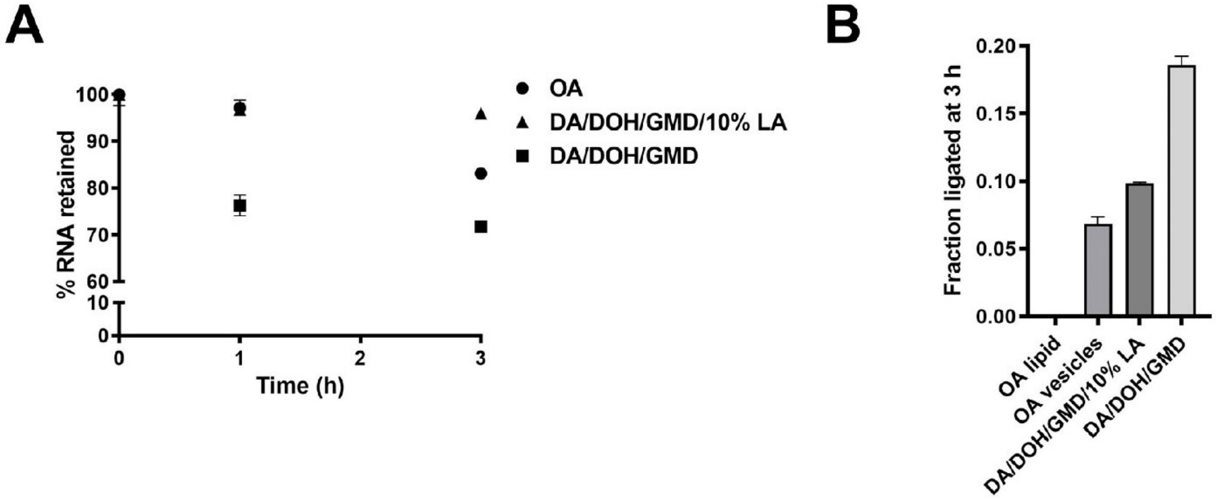


**Figure S7. Ribozyme retention and ligation within vesicles composed of longer lipid chains. A.** Ribozyme leakage from vesicles made of DA/DOH/GMD (4:1:1), DA/DOH/GMD (4:1:1) doped with 10% lauric acid (LA, C12:0), and oleic acid (OA, 18:1) was monitored over 3 h in the presence of 3 mM $Mg^{2+}$. 95% retention was observed for LA-doped DA/DOH/GMD vesicles, while OA and DA/DOH/GMD vesicles showed 83% and 70% retention, respectively. **B.** RNA-catalyzed RNA ligation yields within OA, DA/DOH/GMD/10% LA and DA/DOH/GMD were ~7%, ~10%, and ~19%, respectively after 3 h in the presence of 3 mM $Mg^{2+}$. Ligation reactions containing 2 µM FAM-labeled ribozyme (RS8_trunc), 3 µM RNA template, 4 µM 2AI-activated substrate, and 100 mM Tris-HCl (pH 8.0).

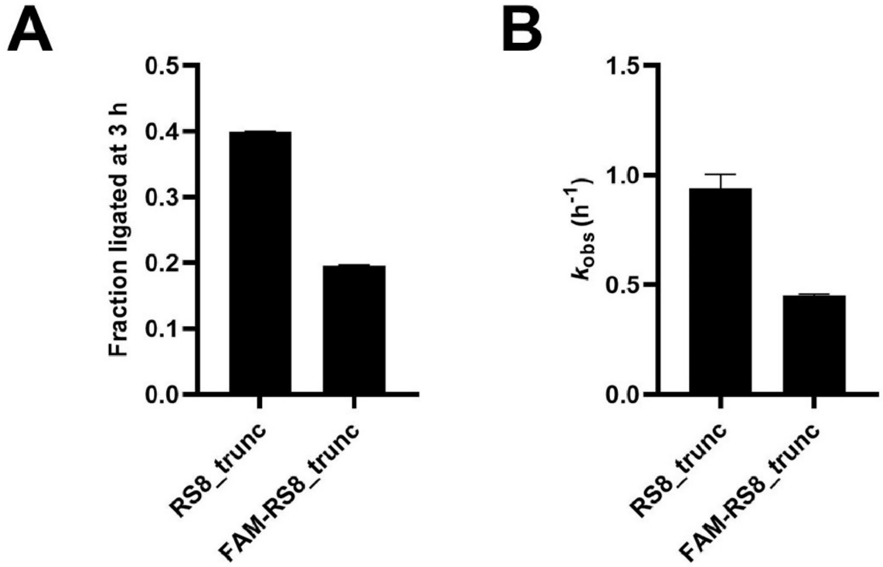


**Figure S8. FAM-labeled RS8_trunc ribozyme is less efficient than the corresponding unlabeled ribozyme. A.** Ligation yield afforded by unlabeled RS8_trunc was twice that by FAM-labeled RS8_trunc. **B.** Unlabeled RS8_trunc was two-fold faster than FAM-labeled RS8_trunc. Ligation reactions containing 2 µM FAM-labeled ribozyme (RS8_trunc), 3 µM RNA template, 4 µM 2AI-activated substrate, 100 mM Tris-HCl (pH 8.0), and 3 mM $Mg^{2+}$.

### Author Contributions

S.D., S.J.Z. and J.W.S. planned the experiments. S.D. and S.J.Z. and M.P.S carried out the experiments and analyzed the data. J.W.S. obtained the funding and supervised the project. S.D., S.J.Z. and J.W.S wrote the manuscript. S.D. and S.J.Z. contributed equally.
